# Label-free three-dimensional analyses of live cells with deep-learning-based segmentation exploiting refractive index distributions

**DOI:** 10.1101/2021.05.23.445351

**Authors:** Jinho Choi, Hye-Jin Kim, Gyuhyeon Sim, Sumin Lee, Wei Sun Park, Jun Hyung Park, Ha-Young Kang, Moosung Lee, Won Do Heo, Jaegul Choo, Hyunseok Min, YongKeun Park

**Author notes:** These authors contributed equally to this work (Jinho Choi, Hye-Jin Kim).

## Abstract

Visualisations and analyses of cellular and subcellular organelles in biological cells is crucial for the study of cell biology. However, existing imaging methods require the use of exogenous labelling agents, which prevents the long-time assessments of live cells in their native states. Here we propose and experimentally demonstrate three-dimensional segmentation of subcellular organelles in unlabelled live cells, exploiting a 3D U-Net-based architecture. We present the high-precision three-dimensional segmentation of cell membrane, nucleus membrane, nucleoli, and lipid droplets of various cell types. Time-lapse analyses of dynamics of activated immune cells are also analysed using label-free segmentation.

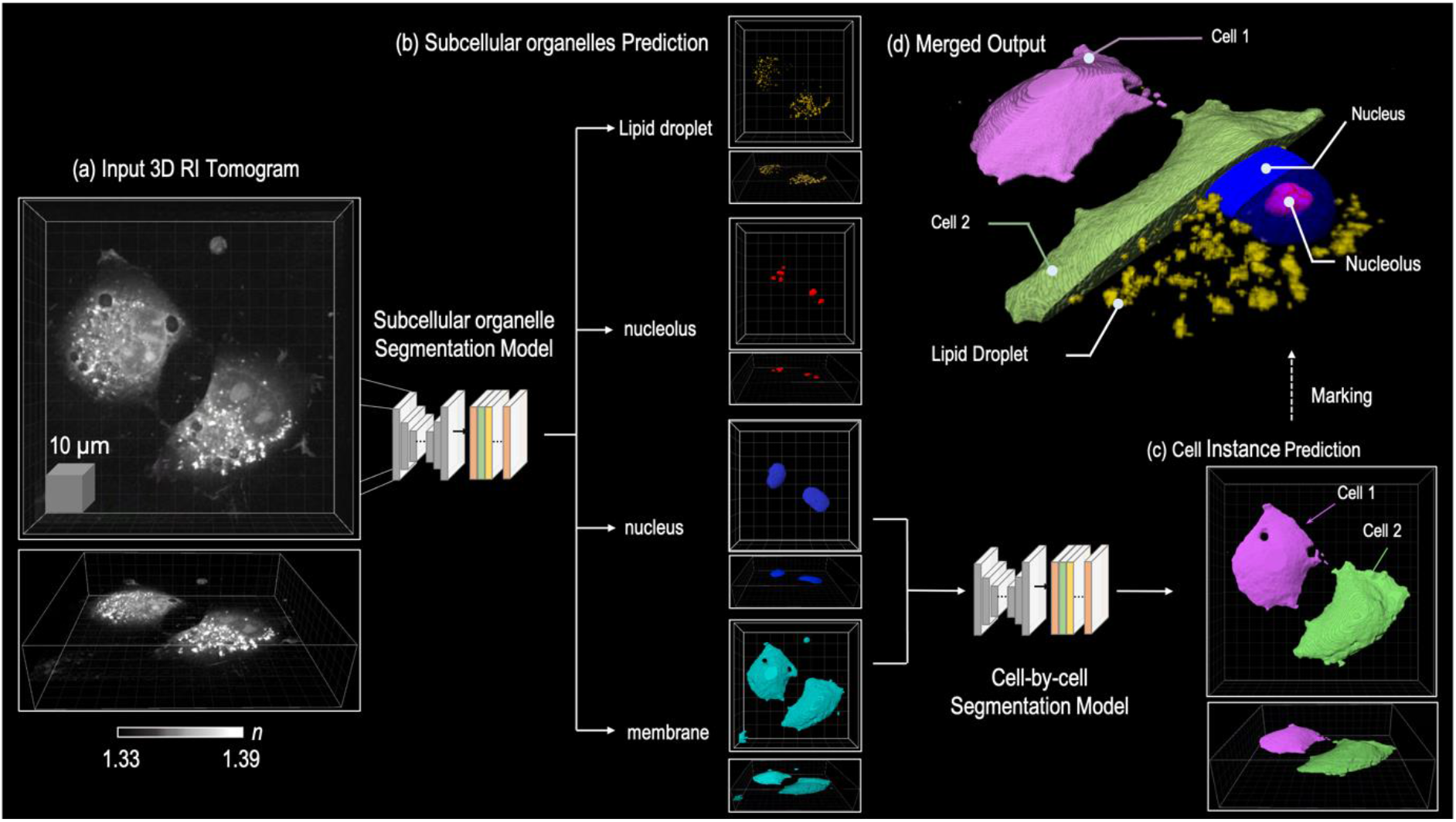

## Introduction

There is a high demand for the quantification of the morphological dynamics in a live cell and its subcellular organelles among numerous research topics in quantitative cell biology^1, 2^. Recent advances in microscopic techniques have created a new era for image-based cell volume quantification^3–5^. Fluorescence-based confocal imaging is the most popular for live-cell quantification, offering high flexibility of organelle markers and correlated fluorophores.

Quantitative phase imaging (QPI) is a powerful method to observe the morphology of a live specimen without any perturbation; this includes dye staining or fluorescence protein expression^6^. Recently developed three dimensional (3D) QPI techniques provide the 3D refractive index (RI) distributions, containing quantitative information on the concentration of a material, and have been exploited in various applications including biomolecular condensates^7^, biotechnology^8^, microbiology^9^, and cell biology^10^. Although the 3D QPI image can provide the physical properties corresponding to each voxel, a universal and versatile segmentation method is required to simultaneously monitor quantitative dynamics in a whole cell and its organelles. To this end, there is a need for techniques to discriminate specific organelles within a cell and discriminate each cell unit from its neighbouring cells.

To provide such a cell segmentation mask in 3D QPI, previous works have widely used conventional approaches such as the threshold-based Otsu segmentation, transforming a 3D image to a two-dimensional (2D) image by maximum intensity projection, and filtering in 3D volume organelle segmentation^11–13^. However, this algorithm may rarely be applied to the organelle segmentation of QPI images due to the lack of organelle specificity from the intensity and the low variation of numerical contrast. As the RI is an intrinsic value determined exclusively by the concentration of a certain material, the RI range can easily overlap among different compartments within a cell.

In recent years, machine learning techniques based on cell and organelle morphology have been adopted to overcome these problems in 3D QPI imagery. In particular, deep-learning approaches based on a large amount of data rather than specific features have been utilised, including nucleus segmentation^14^, spermatozoon segmentation^15^, and lipid droplet segmentation^16^. These semantic segmentation methods should use single-cell images for cell analysis due to the absence of a method to distinguish individual cell units. To overcome this limitation of semantic segmentation in cellular studies, several studies have proposed cell-by-cell segmentation to track the immunological synapse of immune cells^17^ or analyse sperm cells^15^. However, as these methods mainly focus on the segmentation of specific cell types or organelles, their applicability is limited in a few analyses. To be used in various applications, it is necessary to develop a robust model that accurately segments individual cells and organelles among numerous cell types.

This study presents a universal framework for the label-free, quantitative analysis of live cells; this study has three major contributions. First, to simultaneously monitor the quantitative dynamics of whole cells and organelles, an automatic segmentation framework using deep learning and cell characteristics in 3D QPI images was proposed. The proposed automated segmentation framework consists of a “multi-organelle segmentation” model that segments multiple organelles within a cell and a “cell-by-cell segmentation” model that distinguishes individual cells from neighbouring cells. Second, we verified that this model has spatio-temporal robustness among numerous adherent and suspension cell lines, popular among biologists. The proposed framework did not target a specific organelle, but rather it learned the relationship of organelles within a cell, by considering multiple organelles simultaneously. As such, it showed stable performance even within a variety of cells not used for learning. In particular, the cell-by-cell segmentation model operated in various cell lines without being limited to specific cells, based on the cell membrane and nuclear information. Finally, we demonstrate quantitative analyses of RAW 264.7 cells utilising morphological and biochemical properties by exploiting the linear correlation among RI, protein density, and the proposed segmentation models. The results suggest that the proposed method offers a new analytical approach for automatic cell studies.

## Results

### Deep-learning-based multi-organelle and cell-by-cell segmentation model

The proposed analysis process for live cells was primarily composed of two processes. First, we generated segmentation masks for individual cells and their organelles. Then, we obtained the morphological and physical properties (e.g., volume, surface area, and concentration), using each created segmentation mask and its RI values.

To this end, we utilised data-driven, deep-learning techniques. Specifically, we used two different 3D convolutional neural networks: one for the multi-organelle segmentation model, and the other for the cell-by-cell segmentation model, as depicted in Figure 1. The multi-organelle segmentation model predicts the segmentation mask of four organelles from the input 3D RI tomogram (Figure 1a); the nucleus, nucleolus, plasma membrane, and lipid droplet (Figure 1b). We selected these four organelles because they are commonly used in cell analysis. By learning various tasks simultaneously, the model learns the characteristics of individual organelles and their relationship with each other. This multi-task learning prevents overfitting^18^ and significantly reduces computation time, compared to training each task separately.

**Figure 1.**
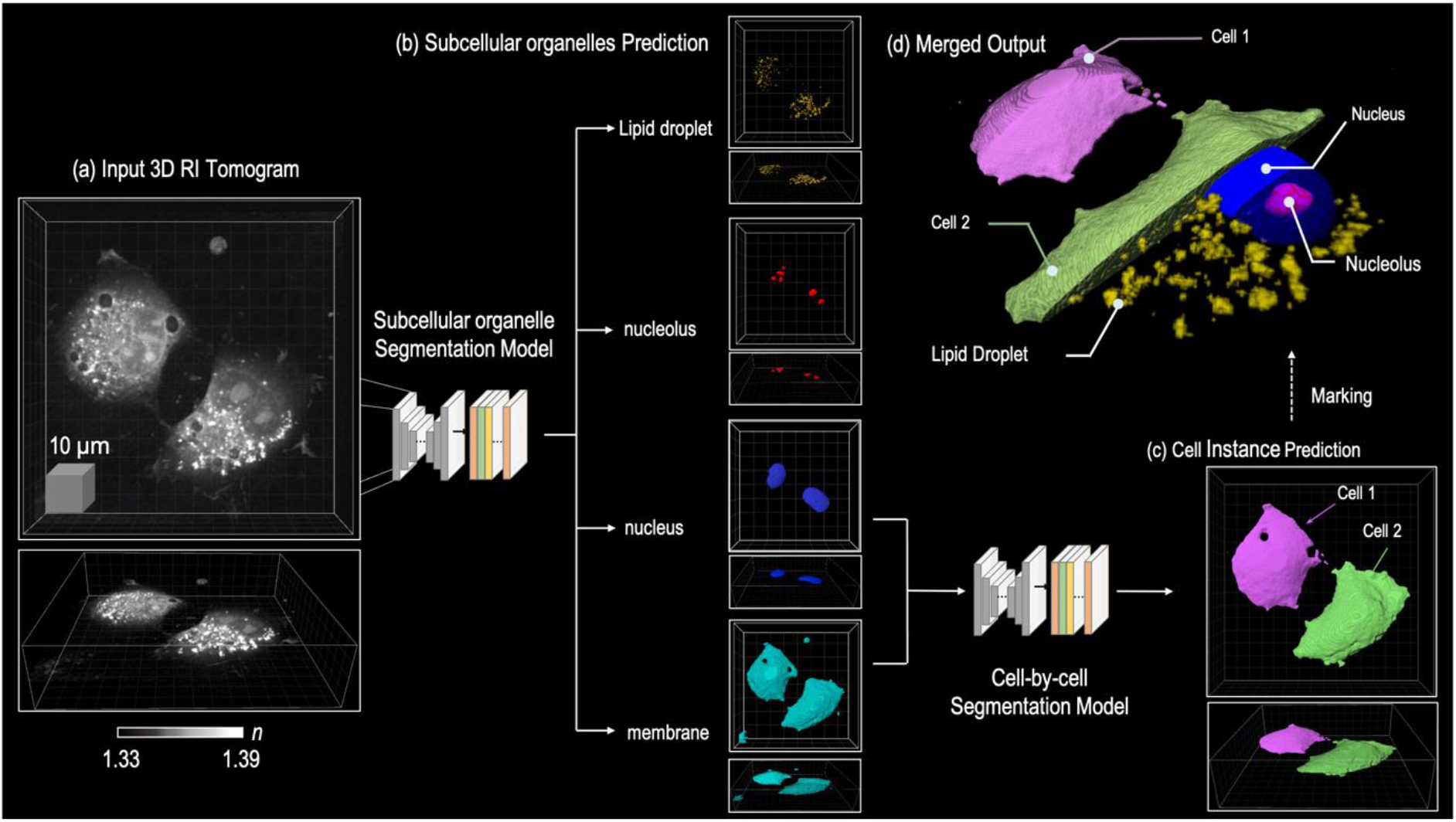
Overview of the multi-organelle segmentation and cell-by-cell segmentation models. **(a)** From the input 3D Tomogram, multi-organelle segmentation model predicts segmentation masks of four subcellular organelles: the membrane, nucleus, nucleolus, and lipid droplets. **(b)** Cell-by-cell segmentation model predicts instance masks by utilising the predicted membrane and nucleus masks. **(c)** Examples of predicted segmentation masks.

The cell-by-cell segmentation model divides the membrane mask of the entire cell into the mask of each cell. The model uses the nuclear and membrane masks obtained from the multi-organelle segmentation model results (Figure 1c). Assuming that each cell has at least one nucleus, a nucleus mask was used as the seed to separate individual cells. The membrane mask was used to distinguish regions between non-cell and cell areas. The details of the model are described in the **Online Methods** section.

We measured 129 3D QPI images of live NIH3T3 cells to train and evaluate the segmentation models described above. To generate ground-truth masks of the collected 3D QPI imagery, three expert biologists manually annotated the masks of individual cells and their four organelles using the open-source software called Insight Segmentation and Registration Toolkit (ITK)-Snap^19^. Thereafter, we split collected data into 105 training and 24 evaluation datasets. Additionally, we used four different cell lines (A549, MDA-MB-231, HeLa, and RAW 264.7) that were not used for training to confirm model generalizability, whereby each cell line was extracted from a different organ. A549 is a human lung carcinoma cell line, MDA-MB-231 is a human breast adenocarcinoma cell line, HeLa is a human cervical adenocarcinoma cell line, and RAW 264.7, a mouse macrophage cell line; nine 3D QPI images were captured for each cell line. These cell lines were used because these cells represent the characteristics of other organs and because they have widely used cell lines in biological laboratories. The preparation of cell lines and the process of generating QPI data are detailed in the **Online Methods** section.

The Dice score was used to quantitatively measure the segmentation performance of the model. This is the most frequently used metric in image segmentation, quantifying the similarity between the ground truth and prediction masks (Eq. 3). The average Dice score of cell instance segmentation for the five cells was 0.758, while the average Dice score of membrane segmentation was 0.831. For the NIH3T3 and RAW 264.7 cells, the difference between the Dice score of membrane segmentation and that of cell-by-cell segmentation was marginal. However, for the remaining cells, the Dice score of cell-by-cell segmentation was slightly lower than that of membrane segmentation. As these cell lines were characterised by confluent growth, the resulting cell-by-cell segmentation tasks were very difficult. Likewise, the Dice scores of the nucleus segmentation for the NIH3T3 and Raw 264.7 cells were higher than the remaining cells. The variation in the Dice score for nucleolus segmentation was relatively small; this is because the RI of the nucleolus is similar between cells. The Dice score of the lipid droplet segmentation was far lower than other organelles, as the volume of the lipid droplet was relatively small compared to the other subcellular organelles. A small portion of false-positive and false-negative predictions may significantly reduce the Dice score when the total volume of the mask is small.

Next, we conducted a qualitative assessment using experts (Figure 2). The cell-by-cell segmentation model performed well in the A549, MDA-MB-231, HeLa, and RAW 264.7 cell lines, which were not used for training. Each of the five cell lines had different shape characteristics in the subcellular organelles. NIH3T3 has a small apparent nucleus, a small number of nucleoli, and a long and overlapping membrane structure. The A549 cells also had an evident trim nucleus, although they possessed one or two large nucleoli, and had thin membrane with large lipid droplets. The HeLa cells have a large prominent nuclear membrane and a small number of nucleoli in the nucleus, with a thicker membrane than A549 cells. The MDA-MB-231 cells were possessed, but it has one or two large nucleoli. The morphology of RAW 264.7 cells completely differed from that of the four other cell lines; the size of RAW 264.7 cells was smaller than that of the other five cell lines; moreover, the RAW 264.7 cells had a spiky circular membrane. The size of the nucleus was sufficiently large to make up most of the cells. We applied the model to cell lines with different characteristics and observed that it worked very well with various cell lines. Additionally, the masks produced by 3D cell segmentation showed better morphological features of cells and subcellular organelles (Figure 3).

**Figure 2.**
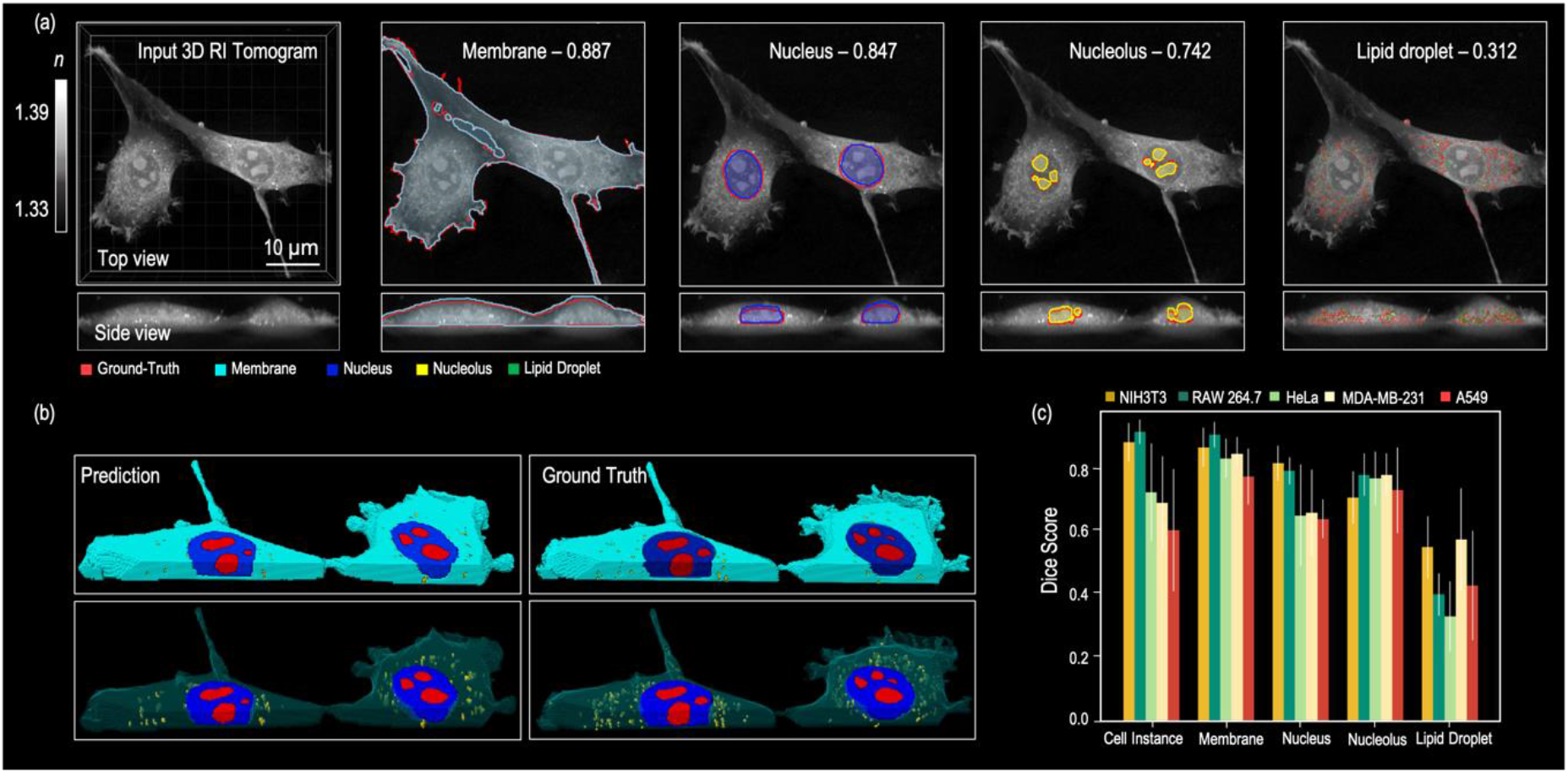
Quantitative results and examples of Dice scores with corresponding prediction and ground-truth masks. **(a)** The numbers indicate Dice scores between prediction and ground-truth masks; the 2D slices of prediction masks and ground-truth masks are overlaid. **(b)** 3D masks of predictions and ground-truth. **(c)** Quantitative results on the NIH3T3, RAW 264.7, HeLa, MDA-MB-231, and A549 cells.

**Figure 3.**
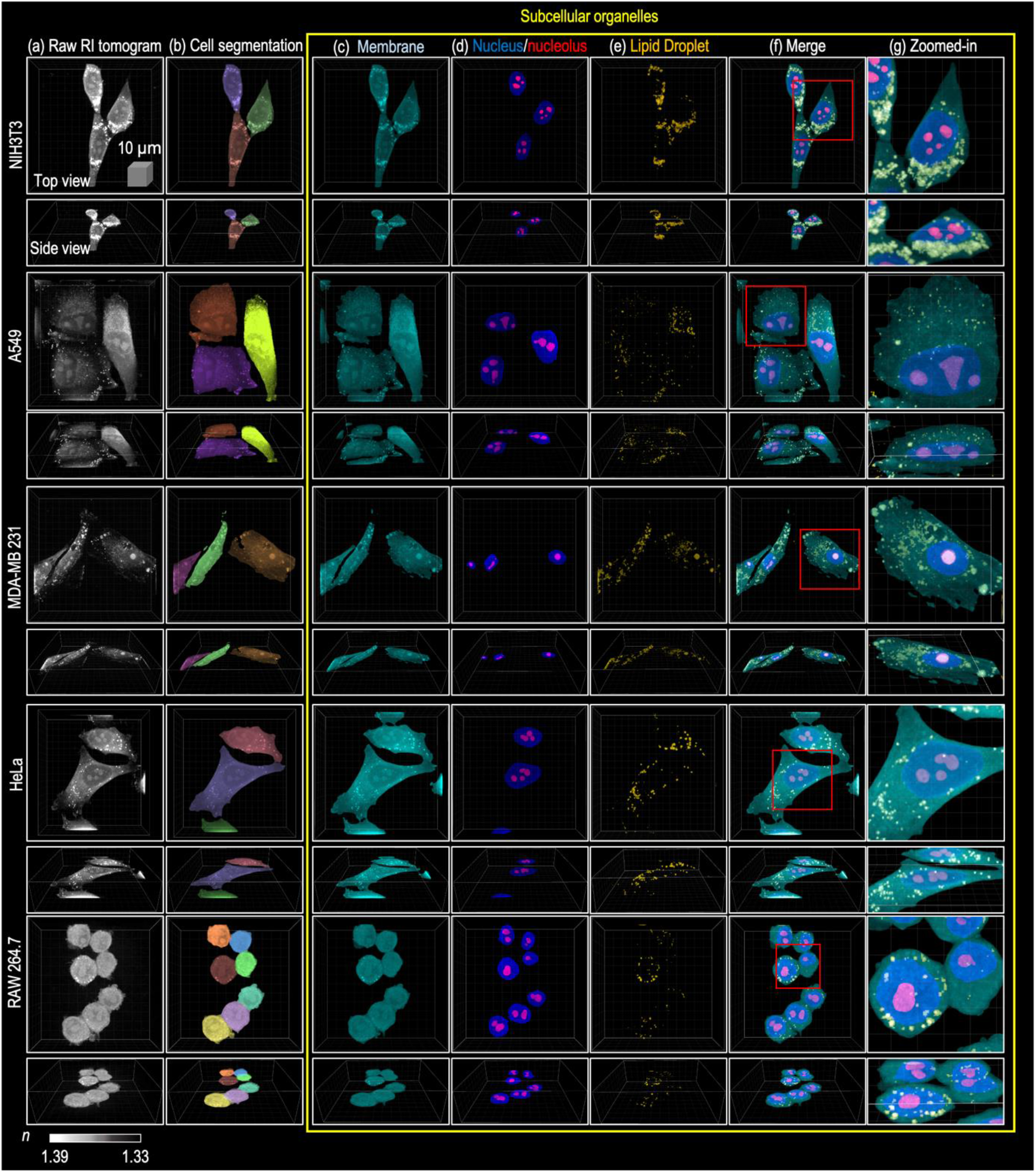
Examples of qualitative results for the NIH3T3, A549, MDA-MB-231, HeLa, and RAW 264.7 cells. **(a)** 3D Input Tomogram. **(b)** Segmentation masks of cell instances. **(c)** Segmentation masks of membrane. **(d)** Segmentation masks of nucleus and nucleolus. **(e)** Segmentation masks of lipid droplet. **(f)** Merged segmentation masks of four subcellular organelles. **(g)** Zoomed-in patches of inputs.

### Quantitative cell analysis using segmented masks

To analyse live cells, we utilised volume, surface area, and concentration. Volume was computed by multiplying the total number of voxels in the segmentation masks and the volume of the 3D QPI image. To compute surface area, we constructed a triangular mesh from the 3D segmentation masks. Concentration was calculated as per Equation (1):

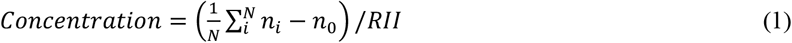

where *N* is the index of the set of the segmentation mask; *n_i_* is the RI of the voxel; *n*_0_ is the RI of the surrounding media (1.3337); the RII is a constant set as 0.135 for the lipid droplet and as 0.19 for the remaining organelles.

Macrophages are white blood cells that play an essential role in the innate immune system. Macrophages phagocytose bacteria, secreting proinflammatory and antimicrobial mediators^20^. Macrophages induce the transcription of genes that encode proinflammatory regulators, such as the transcription factor activator protein 1 (AP-1), and the c-Jun N-terminal kinase (JNK). This is achieved through exposure to environmental stimuli, known as macrophage activation^21^. Activated macrophages are also known to change phenotypes, although the details are not well defined. Activated macrophages have two different phenotypes depending on exposure stimuli: M1 and M2. Macrophage polarisation causes different behaviours in the immune system, and alters cell shape differently^22^. Although the critical relationship between cell shape and the function of macrophages has been studied, there is little research on the quantitative analysis of shape changes in whole cells and subcellular organelles. We hypothesised that macrophage activation would affect gene and protein expression, and the morphology and physical characteristics of cells and subcellular organelles. To achieve this, a mouse macrophage cell line, RAW 264.7, was used as a model, and macrophage activation was induced through treatment with bacterial lipopolysaccharides (LPS).

Without requiring secondary assays, 3D QPI was able to directly observe and calculate morphological and biochemical changes in activated RAW 264.7 cells. The 3D RI tomogram of LPS-treated RAW 264.7 cells showed that activated cells undergo dramatic morphological changes within 24 h (Supplementary Figure 2). The control RAW 264.7 cells were characterised by the circular shape of a typical macrophage cell, slightly attached to the bottom of the culture dish. However, 8 h after LPS treatment, these RAW 264.7 cells began expanding in volume and attached themselves to the bottom of the dish. At 24 h, the attached cells formed lamellipodia and granules in the cytosol (Supplementary Figure 2). We attempted to track and calculate the changing parameters of RAW 264.7 cells for 8.5 h during the activation process as these RAW 264.7 cells dynamically altered their shape during the initial response to LPS treatment.

The 3D RI tomogram of LPS-treated RAW 264.7 and untreated control cells were acquired every 30 min for up to 8.5 h on a 3D QPI microscope in label-free states. The 3D cell segmentation was conducted with every time-lapse image, generating subcellular organelle masks (Figure 4a). The generated masks represent the changing phenotype of the activating macrophage process. The membrane masks perfectly represented the spreading morphology of activated macrophages overtime in 3D. Figure 4a shows that the verified volume of activated macrophages had become bigger and wider through the membrane mask. In addition to the *xy* slices, the *yz*, and *xz* slices enabled easy identification of the increased volume of the cell membrane through the generated mask. The subcellular organelle masks of the nucleus and nucleolus and lipid droplet were acquired from label-free holographic imagery. Although the size of the nucleus appears to grow along the cell membrane, retention trends were observed in the nucleolus compared to the membrane and the nucleus. The most noticeable changes were observed in lipid droplets; during inflammation, when macrophages recognise inflammatory stimuli such as bacterial LPS, they induce the accumulation of cholesterol esters and triglycerides in their own body^23^. The model was able to precisely separate lipid droplets, and these represent the increasing size of the fat and lipid droplets, as shown in previous studies. At 0 min, the number and size of lipid droplets were small, while 8.5 h after LPS exposure, the size and number of lipid droplets had increased. Figure 4b presents the mean changes in the RI value for the RAW 264.7 cells during the LPS-induced activation process. We measured the protein concentration, surface area, and volume of individual masks from the RI value, tracking these parameters overtime. Consistent with the cell spreading observed in Figure 4a and Supplementary Figure 2, the volume and surface area of RAW 264.7 cells significantly increased after 8 h activation. The mean RI of the RAW 264.7 cell membranes commenced decreasing immediately after LPS treatment, and continued to decrease steadily after 8.5 h of time-lapse measurements. Although the protein concentration of the membrane decreased, the surface area and volume of the membrane increased gradually for 8.5 h (Figure 4a). The nucleus, similar to the membrane, tended to decrease RI and protein concentrations. While the volume and surface area of the nucleus increased, it was confirmed that the volume and area of the nucleolus slightly increased at the beginning, and was maintained for 8.5 h. For the lipid droplets with very high RI compared to other subcellular organelles (e.g., the nucleus and nucleolus), it was observed that the RI was maintained during the activation process for 8.5 h.

**Figure 4.**
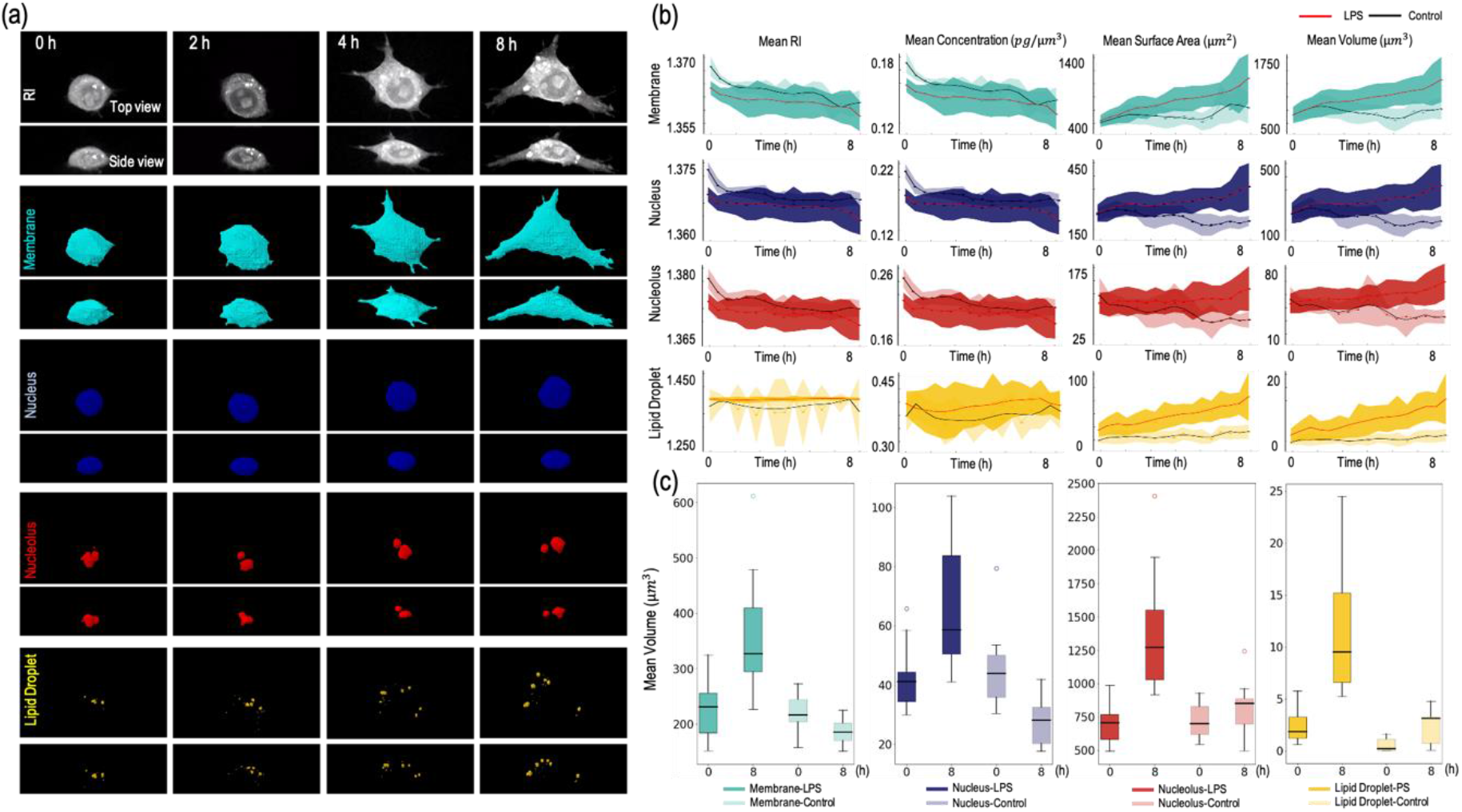
Application to time-lapse imagery of RAW 264.7 cells. **(a)** Horizontal plane and coronal plane view of RAW 264.7 cells overtime (0, 2, 4, 8 h). Each panel represents the RI tomogram and segmentation masks of the membrane, nucleus, nucleolus, and lipid droplet. **(b)** The black line and the light shades show the control, LPS-untreated RAW 264.7 cells (n=9), while the red line and the dark shades show LPS-treated, activating RAW 264.7 cells (n=12). The mean RI and mean protein concentration decreased by activation, while mean surface area and mean volume increased during activation. **(c)** Box plots representing the minimum to maximum and showing all points of the mean volume of subcellular organelles in the RAW 264.7 cells.

The surface area and volume of lipid droplets increased during macrophage activation compared to the control cells, similar to the membrane and nucleus. These observations suggest that LPS-induced changes in macrophages occur phenotypically, and manifest in physical changes. We compared the individual masks of subcellular organelles from the start point to end point of time-lapse data (Figure 4c). The mean RI of each organelle decreased rapidly for 8.5 h, although the volume and surface area increased. The results indicate that RAW 264.7 cells were increasing in size while losing their concentration during the activation process. This occurred throughout the whole cell and in subcellular organelles, including the nucleus, nucleolus, and lipid droplets.

## Discussion and Conclusion

The results show that the proposed framework, combining 3D QPI with deep neural network-based segmentation models, enables label-free 3D live cell analysis in an automated manner. The proposed framework predicts the segmentation mask of organelles within individual cells and uses RI to provide physical and morphological information on cells and organelles. In particular, to automatically segment each cell, we assumed that each cell had at least one nucleus. We used the predicted nucleus segmentation mask of the cell as seed information to distinguish each cell. The proposed framework did not target specific organelles. Rather, it predicted multiple organs concurrently. By training the model to segment several organelles simultaneously, the model learned the relationship of organelles and showed stable performance even in various cell lines not used for training. We also demonstrated that existing biological knowledge may be confirmed through the proposed framework by automatically tracking and observing cell dynamics in time-lapse data.

To the best of our knowledge, this work is the first of its type to analyse various 3D cell organelles simultaneously and automatically at an instance level. An immediate future research priority lies in further improving model performance. Introducing Bayesian neural networks^23^ and stable learning methodologies using uncertainty prediction^24, 25^ should be considered for a more robust model. In future research, this framework should be expanded to refine the predicted segmentation masks utilising user interactions. When combined with cell lineage tracking technology, changes in the temporal differentiation process of cells may be automatically monitored. In addition, it can support real-time analysis by improving the efficiency of networks to process large 3D imagery.

## Online Methods

### Deep-Learning Models

#### Subcellular organelle segmentation

The multi-organelle segmentation model uses a 3D RI tomogram image as input and predicts the binary mask of four organelles: the nucleus, nucleolus, membrane, and lipid droplets (Supplementary Figure 1). We train the model to predict the mask of four different organelles simultaneously, as opposed to training an independent model for each organelle. This approach, known as multi-task learning, improves the overall performance of multiple tasks^18^. In addition, the model training time was drastically reduced by using only a single model.

3D RI tomogram images have high resolutions, varying from 100×600×600 to 260×860×860 voxels. As such, training the subcellular organelle segmentation model using the entire volume requires a huge graphics processing unit (GPU) memory. For this reason, during the training phase, we initially resized the input 3D RI tomogram to 128×512×512 voxels. Then, we randomly sampled patches of 64× 128× 128 from the resized 3D RI tomogram and utilised these as inputs. The model predicted the probability map for each subcellular organelle producing an identical resolution to the input patch.

During the inference phase, we resized the input to 128×512×512 voxels and applied a 64-size symmetric padding. We cropped the input image from the centre along the *z* axis with the size of 64. Then, we uniformly generated patches of 64× 128× 128 with a stride of 128 and obtained probability maps for each patch. We reconstructed the predicted patches into the entire volume of the image by stitching patches using a spline kernel. Following this, we removed the padding area from the stitched probability maps and restored them to the original resolution. Finally, we obtained the segmentation mask by binarising the probability map using a threshold of 0.5.

#### Cell-by-cell segmentation

To predict the instance masks of cells, the cell-by-cell segmentation model utilises segmentation masks of the nuclei and the membrane predicted by the subcellular organelle segmentation model. The nuclei mask was used as prior information regarding the number of cells and their approximate location. The membrane mask prevented the model from predicting the background area as the cell. The model initially separated the predicted nuclei mask into the instance masks of the nucleus, *n_i_* ∈ *R^D×H×W^*, where i ∈ {1,…, k} indicates the index of nuclei; and *k* is the total number of nuclei. As the nuclei of cells were separated, the instance masks of the nucleus were simply obtained using a connected component algorithm. Then, we selected the *i*^th^ nucleus instance mask, *n_i_*, and considered this a positive map, *pos_i_*; the remaining nuclei instances were considered a negative map, *neg_i_*. We concatenated the membrane mask, *m*, and *pos_i_* and *neg_l_* to *I*, predicting the instance probability map (x_i_) of the cell that includes the selected nucleus instance (n_i_).

During the training phase, we randomly selected one nucleus instance and trained the model to predict the instance mask of the selected cell. During the inference phase, the model repeated this process for each nucleus and finally obtained the instance mask by assigning the index of the nucleus that has the highest probability: M = *argmax*_*i*=1,… *N*_*x_i_*, where *N* is the number of nuclei. We considered a voxel as the background if the highest probability was lower than the 0.5 threshold.

In contrast to the subcellular organelle segmentation task, the patching strategy was not applicable to the cell-by-cell segmentation task as the whole-cell shape was critical when predicting the instance mask. Therefore, we downsized inputs to 128× 128 in the *x* and the *y* axes, and cropped the resized inputs along the *z* axis to the size of 64. Then, we restored the predicted instance mask to the original size of the inputs.

#### Network architecture and training details

The 3D U-Net-based architecture was adopted^26^, and this was demonstrated to have impressive performance in biomedical image segmentation tasks, as per these models. Specifically, we employed the Scalable Neural architecture search (ScNas)^27^, which automatically designs the architecture optimised for 3D cell image segmentation. ScNas identifies network parameters and micro-level architectures by utilising a stochastic sampling algorithm. Similar to the U-Net, the constructed networks were composed of encoder and decoder cells; encoder cells extract feature maps at multiple scales by gradually downscaling resolution, while decoder cells up-sample the extracted feature maps to the original resolution and classify the label of voxels. Each cell consists of repeated stacks of 3D convolutional layers, a rectified linear unit (ReLU), pooling operations, and normalisation layers; we utilised the constructed networks for subcellular organelle and cell-by-cell segmentations.

We selected the activation and normalisation functions of ScNas as Leaky-ReLU^28^ and instance normalisation^29^, respectively. The size of the initial feature map, the number of layers, and the feature map multiplier were set to 12, 8, and 3, respectively. Hyper-parameters of the network were adjusted using a grid search algorithm. The models were implemented in Python 3.7, using the PyTroch 1.4 framework on an 8 V100-32G GPU machine.

Several data augmentation strategies were applied, such as random flipping, cropping, and rotation. In particular, the input image was rescaled from 0.5 to 2 to handle varying resolutions of 3D RI tomography. We utilised the Adam^30^ optimiser with the learning rate of 0.001, and reduced the learning rate by the factor of 5 if there was no improvement in the validation metric for 30. To train the models, we combined the Dice loss and the binary cross entropy (BCE) loss; this is defined as:

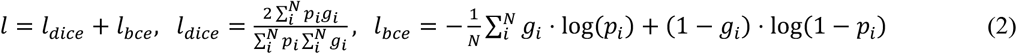

where *N* is the number of voxels; and *p_i_* and *g_i_* indicate the predicted probability and the ground-truth label of the *i*^th^ voxel, respectively. For the multi-organelle segmentation task, we trained the model to conduct multiple tasks simultaneously; thus, the loss for each subcellular organelle was calculated and the sum of all losses was determined. For cell-by-cell segmentation, we simply computed the loss between a selected instance mask and its prediction.

#### Metrics

For quantitative evaluation, we adopted a Dice coefficient that measures the similarity between the predicted mask and the corresponding ground-truth mask; this coefficient is defined as:

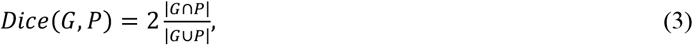

where *G* is the ground-truth mask; and *P* is the predicted mask. For the cell-by-cell segmentation task, the Dice coefficient score, *Dice*(*P_i_, G_j_*). was determined for all pairs of instance masks associated with the prediction and the ground-truth. Then, we applied the Hungarian algorithm^31^ to assign the prediction (*P_i_*) to ground-truth (*G_j_*), which had the highest Dice score.

#### Cell line and cell culture

The NIH3T3, A549, HeLa, and RAW 264.7 cell lines were purchased from the Korean Cell Line Bank (KCLB, Seoul, Korea) and cultured in Dulbecco’s modified Eagle’s medium (DMEM; high glucose) (Hyclone, SH30243) supplemented with 10% (vol/vol) fetal bovine serum (FBS; Hyclone, SH30084) and 1% antibiotic-antimycotic solution (Thermo Fisher Scientific). The MDA-MB-231 cells were purchased from the Korean Cell Line Bank (KCLB, Seoul, Korea) and maintained in RPMI-1640 medium (Hyclone, SH30027) supplemented with 10% (vol/vol) FBS (Hyclone, SH30084) and antibiotic-antimycotic solution (Thermo Fisher Scientific).

#### LPS treatment in RAW 264.7 cells and plating on TomoDish

Precisely, 30 μL of LPS (100 μg/mL stock, List Biological Laboratories) from *Escherichia coli* was added to 3 mL of Dulbecco’s Modified Eagle Medium supplemented with 10% FBS and 1% antibiotic-antimycotic; 3.0× 10^5^ RAW 264.7 cells, a mouse macrophage cell line, were counted and added to a 15 mL tube, and cells were centrifuged at 100 × g for 5 min to collect cell pellets. The supernatant medium was removed using suction, and the cell pellet was gently resuspended in 3 mL of LPS-containing medium. Then 3 mL of medium with RAW 264.7 cells were moved to the TomoDish (Tomocube, Inc.).

#### The 3D QPI

The 3D RI images of cells were obtained using a commercial holotomography (HT-2H, Tomocube Inc., Republic of Korea), based on Mach-Zehnder interferometry equipped with a digital micromirror device (DMD). A coherent monochromatic laser (= 532 nm) was divided into two paths, a reference and a sample beam, using a 2×2 single-mode fibre coupler. A 3D RI tomogram was reconstructed from multiple 2D holographic images acquired from 49 illumination conditions, a normal incidence, and 48 azimuthally symmetric directions with a polar angle (64.5°). The DMD was used to control the angle of the illumination beam impinging on the sample^32^. The diffracted beams from the sample were collected using a high numerical aperture (NA) objective lens (NA=1.2, UPLSAP 60XW, Olympus). To compensate the missing cone issue due to the limited NA, a regularization algorithm based on non-negativity was used^33^. The off-axis hologram was recorded using a complementary metal oxide semiconductor image sensor (FL3-U3-13Y3MC, FLIR Systems). The visualisation of the 3D RI maps and its correlative 3D fluorescence signal with red pseudo-colour was carried out using commercial software (TomoStudio™, Tomocube Inc.). The details on the principle and reconstruction algorithms can be found elsewhere^34, 35^.

#### Time-lapse imaging using a holotomography microscope

Prior to 3D QPI imaging, the HT-2H (Tomocube, Inc.) was turned on to warm up the laser for at least 30 min. Additionally, the carbon dioxide (CO2) gas mixer and temperature controller were turned on to maintain a temperature of 37°C and an atmosphere with 5% CO2 in the TomoChamber (Tomocube, Inc.). The water reservoir of the TomoChamber (Tomocube, Inc.) was filled with autoclaved distilled water to maintain humidity during imaging.

Immediately after LPS treatment, RAW 264.7 cells containing TomoDish were mounted on the TomoChamber, and this chamber was then gently mounted on the HT-2H stage. Then, the HT-2H was calibrated, and the acquisition tab was used to begin setting up time-lapse imaging. For RAW 264.7 cell time-lapse imaging, 3D QPI images were taken at 15 different positions every 30 min for 17 counts; this was completed in 8.5 h to acquire 3D QPI time-lapse imagery.

## Acknowledgments

This work was supported by KAIST UP program, BK21+ program, Tomocube, National Research Foundation of Korea (2017M3C1A3013923, 2015R1A3A2066550, 2018K000396, 2019R1A2C4070420, 2020R1A2C3014742), and Institute of Information & communications Technology Planning & Evaluation (IITP) grant funded by the Korea government (MSIT) (2021-0-00745, 2019-0-00075).

## Author Contributions

S. Lee, H. Min, and Y.K. Park conceived the idea. J. Choi, H.-J. Kim, G. Sim carried out the measurements and analyses. J. Choi, G. Sim, J. Choo, and H. Min designed, implemented and optimized the deep learning pipeline. H.-J. Kim, S. Lee, W.-S. Park, J. Park, H. Kang designed the experiments, collected data. S. Lee and W.-S. Park established the molecular biology and imaging. All the authors wrote the manuscript. Y.K. Park supervised the work.

## Statement of Competing Interests

J. Choi, H.-J. Kim, S. Lee, W.-S. Park, J. Park, H, Kang, and Y.K. Park have financial interests in Tomocube Inc., a company that commercialises optical diffraction tomogram, and quantitative phase imaging instruments and is one of the sponsors of the work.

