## Supplementary Information for "Label-free three-dimensional analyses of live cells with deep-learning-based segmentation exploiting refractive index distributions"

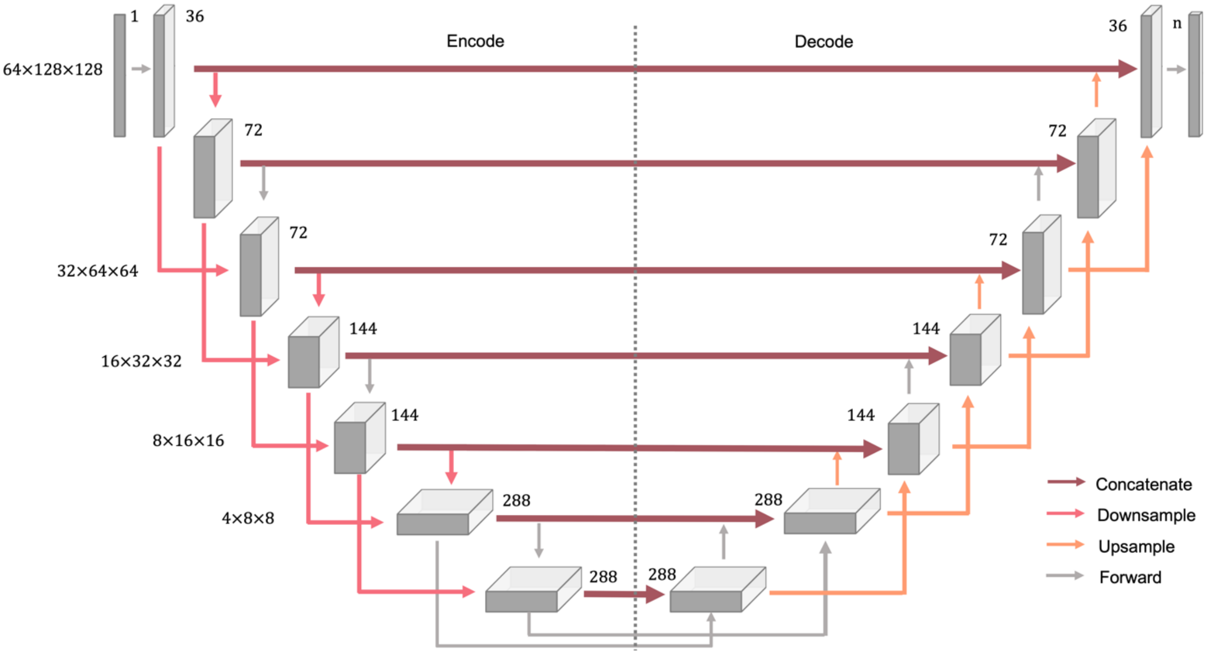


**Supplementary Figure 1. Model Architecture.**

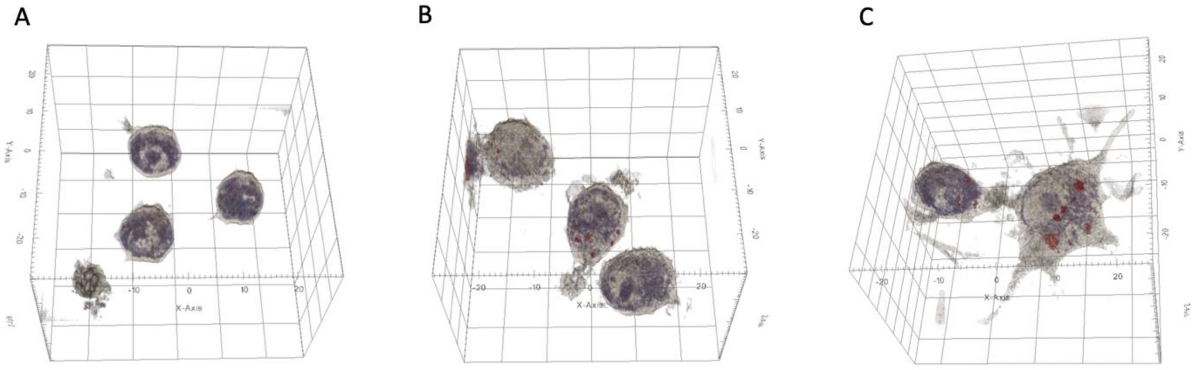


**Supplementary Figure 2. The 3D RI tomogram of LPS-treated RAW 264.7 cells. (a–c)** The 3D rendered isosurface image of 3D RI distribution of **(a)** untreated (start point), **(b)** 8-h LPS-treated, and **(c)** 24-h LPS-treated RAW 264.7 cells.
